# Automated neuron tracking inside moving and deforming animals using deep learning and targeted augmentation

**DOI:** 10.1101/2022.03.15.484536

**Authors:** Core Francisco Park, Mahsa Barzegar Keshteli, Kseniia Korchagina, Ariane Delrocq, Vladislav Susoy, Corinne L. Jones, Aravinthan D. T. Samuel, Sahand Jamal Rahi

## Abstract

Advances in functional brain imaging now allow sustained rapid 3D visualization of large numbers of neurons inside behaving animals. To decode circuit activity, imaged neurons must be individually segmented and tracked. This is particularly challenging when the brain itself moves and deforms inside a flexible body. The field has lacked general methods for solving this problem effectively. To address this need, we developed a method based on a convolutional neural network (CNN) Awith specific enhancements which we apply to freely moving *Caenorhabditis elegans*. For a traditional CNN to track neurons across images of a brain with different postures, the CNN must be trained with ground truth (GT) annotations of similar postures. When these postures are diverse, an adequate number of GT annotations can be prohibitively large to generate manually. We introduce ‘targeted augmentation’, a method to automatically synthesize reliable annotations from a few manual annotations. Our method effectively learns the internal deformations of the brain. The learned deformations are used to synthesize annotations for new postures by deforming the manual annotations of similar postures in GT images. The technique is germane to 3D images, which are generally more difficult to analyze than 2D images. The synthetic annotations, which are added to diversify training datasets, drastically reduce manual annotation and proofreading. Our method is effective both when neurons are represented as individual points or as 3D volumes. We provide a GUI that incorporates targeted augmentation in an end-to-end pipeline, from manual GT annotation of a few images to final proofreading of all images. We apply the method to simultaneously measure activity in the second-layer interneurons in *C. elegans*: RIA, RIB, and RIM, including the RIA neurite. We find that these neurons show rich behaviors, including switching entrainment on and off dynamically when the animal is exposed to periodic odor pulses.

## Introduction

Whole-brain imaging with single-cell resolution is widely used to study the neural circuits for behavior in many organisms including *C. elegans, Drosophila*, zebrafish, and hydra [1–4]. Fast 3D microscopes – light-sheet, spinning disk confocal, light-field, and multifocus microscopes – are expanding the range of brain imaging with fluorescent genetically-encoded calcium indicators to many animals and their behaviors [5–10]. It is often preferable to study brain dynamics in animals as they perform behaviors without restraint: immobilization can change brain activity [11] and many behaviors such as mating or predation only occur in moving animals in natural contexts [12, 13].

Analyzing whole-brain imaging datasets requires solving two problems. One problem is segmentation – the pixels for each neuron must be separated and identified in each image volume. Another problem is tracking – a given neuron must be correctly identified in every image volume. These problems are especially challenging in animals with flexible bodies like nematodes or hydra. Their neurons are small, densely packed, and follow complex trajectories as the brain deforms during behavior. Genetically-encoded labeling of neurons creates additional challenges in imaging and analysis. Conditions for microscopy can vary with different animals and different experiments. Expression patterns of fluorescent reporters can vary from animal to animal. When imaging with dim reporters inside moving animals, neuronal signals may be intermittent and neurons may drop out of view at some time points.

A solution to segmenting and tracking neurons during brain-wide imaging has stringent requirements. In every neuron, every signal at every time-point might contain useful information. Recording circuit activity requires accurate and reliable signal analysis throughout all image volumes. Reliable analysis of image volumes often involves laborious manual annotation. In a recent study of the mating circuit of male *C. elegans*, we required 200 hours to manually annotate 76 neurons in image volumes recorded at 5 Hz in each 10 min experiment [12]. Automated annotation is needed to accelerate the field of brain-wide imaging. Because automated annotation can always generate errors, it must be followed by comprehensive proofreading. However, if automated annotation were sufficiently fast and reliable, the time required to manually proofread an entire dataset could become less than the time required for full manual annotation.

One way to simplify the problem of neuron segmentation is to restrict fluorescent labeling to cell nuclei. In each image volume, cell nuclei form a constellation of non-overlapping and nearly spherical volumes. With their stereotyped appearance, cell nuclei are, in principle, suitable for segmentation by standard methods in image processing. One method involves identifying objects that have a certain size and convex shape along all axes [14]. After objects have been segmented, they need to be tracked across image volumes. One approach to the tracking problem is to seek the optimal alignment of different 3D brain images using methods from point-set registration [15]. These techniques have been applied to *C. elegans* by treating cell nuclei as points in space, and then searching for and registering the local constellation of points surrounding each point in an image volume [16]. However, these methods have not been extended to segmenting and tracking neurons as 3D volumes.

Another approach to tracking neurons is to identify all segmented neurons in each image volume and assign them a unique label computationally. This is possible in animals like *C. elegans* where all neurons have unique identities, and can be done by building probabilistic models of the positions of all neurons using brain atlases that describe a given posture. For each brain image, one calculates the most likely distribution of unique neurons [17–19]. However, this approach is not easily extended to moving, deforming brains, and has only been applied to immobilized animals. [17]

A third way to track neurons is to characterize the collective 3D motion trajectories of neurons over time. In the rapidly deforming body of the hydra, an effective particle motion tracking algorithm has been developed that calculates the most likely set of collective movements of visible neurons [20].

Alternatively, neuron segmentation and tracking during brain-wide imaging can be viewed as problems in pattern recognition, for which deep neural networks are ideally suited. [21] Deep neural networks have been used to perform point-set registration when tracking neurons by training the networks to find the most likely alignment of 3D images. Deep neural networks have also been used to learn the most likely trajectory in collective motion tracking [22, 23]. These approaches facilitated the tracking of neurons, but began with accurately segmented images. In addition to requiring high-quality segmentations, 3DeeCellTracker [22] was tested only on worms that were computationally straightened using additional low-magnification images. fDLC [23] aims to allow deformations, however, it is unclear how the method, which relies on point clouds, would be applied to tracking neurons that are represented as 3D shapes.

We sought a comprehensive pipeline that begins with raw unsegmented brain-wide recordings and ends with proofreading of all tracked and segmented neurons, each neuron represented either as a key-point at its nucleus or as a 3D shape. To be generalizable, we did not want the pipeline to use information beyond the brain images themselves, such as low-magnification images of brain or body posture that are used in some approaches [16, 23]. To do this, we developed a deep learning method that is both significantly faster than full manual annotation, and that functions end-to-end by simultaneously segmenting and tracking neurons in freely moving *C. elegans*.

Brain-wide imaging is subject to substantial experiment-to-experiment variations. To be robust, a convolutional neural network (CNN) must be separately trained using ground-truth data that comes from each experiment. Generating annotated training data for every experiment is labor intensive. We thus sought to minimize the amount of manually annotated data required for each experiment. We developed ‘targeted augmentation’, a means of synthesizing large amounts of annotated training data for the many different shapes and postures of the brain in a given experiment. Our pipeline learns a set of internal deformations of a freely moving *C. elegans*. The pipeline uses these deformations to generate synthetic image volumes and annotations from a small number of original image volumes and their manual annotations. When a CNN is trained with both manual annotations and synthetic annotations for each experiment, its accuracy and reliability dramatically improve.

We further increased the accuracy and reduced the computational burden of the CNN by developing a new architecture to deploy targeted augmentation. Our low error rate for automated annotation substantially reduces the time required for final proofreading. We implement all steps in the pipeline – manual annotation, analysis, and proofreading – in an easy-to-use graphical user interface (GUI). The GUI also implements additional machine learning methods to speed up the initial manual annotation.

By reducing the amount of manually annotated training data required to train a CNN to reliably perform both neuron segmentation and tracking across time, as well as reducing the amount of proofreading required to remove errors, our pipeline achieves a 3x-fold increase in analysis throughput in comparison to full manual annotation for the most challenging brain imaging problem in *C. elegans* to date from ref. [12].

Because our method works both for key-point as well as volumetric segmentation and tracking, we used our pipeline to analyze recordings of freely moving *C. elegans* with labeled neurons that need to be analyzed as 3D volumes, which cannot be easily accomplished with previous methods. Specifically, we investigated the second-layer interneurons in *C. elegans*: RIA, RIB, and RIM. This set of neurons is thought to be important for sensorimotor integration [24, 25] and each has been found to be associated with different functions during *C. elegans* chemotaxis: RIA shows compartmentalized calcium activity [26, 27], in which different segments of the RIA neurite encode dorsal and ventral head movements, respectively, but the soma does not show prominent calcium activity. Thus, RIA has to be segmented and tracked as a 3D volume. RIB activity promotes forward locomotion [2, 28–30]. RIM depolarization extends reversals, while hyperpolarization extends the forward motor state [31]. We asked whether these interneurons also represented olfactory stimuli in addition to their tight link to chemotactic behavioral output. We applied periodic inputs to behaving animals in the form of odor pulses to dissect fundamental network and circuit properties [32–35]. We observed that second-layer interneurons switched between entrained responses to the sensory input or non-entrained activity, indicating switches between states where these interneurons couple or decouple from sensory information.

In summary, our main achievements are 1) a ‘targeted augmentation’ method that reduces the need for manual annotation to create training datasets for a CNN that solves the segmentation and tracking problem in brain imaging, 2) a new CNN architecture that is optimized for this application, 3) a generalizable pipeline, implemented in a graphical user interface (GUI), that is able to track neurons as either key-points or 3D shapes using only information contained in the brain images themselves (no pre-training or additional recordings necessary), and 4) the application of our method to track both the nuclei of the second-layer interneurons RIB and RIM as well as the whole 3D volume of RIA, including its (thin) neurite, showing the complex coupling of these neurons to sensory information. Furthermore, we made the GUI and 4D image datasets for testing and further method development freely available, see ‘Code and data availability’.

## Results

### Whole-brain recording in behaving animals

It is now possible to measure the activity of an entire *C. elegans* brain with cellular resolution during animal behavior using fast 3D imaging systems such as spinning-disk confocal microscopy adapted for multi-color, multi-neuron, real-time tracking [5, 12, 37, 38]. Brain-wide imaging in *C. elegans* is usually performed using transgenic strains with panneuronal expression of a nuclear-localized calcium indicator (e.g., GCaMP6s) and red fluorescent protein (e.g., mNeptune). Alternatively, subsets of neurons may be fluorescently labeled throughout their cytosols to allow recording from cell bodies and neurites. Stable, red fluorescence signals are used to isolate and track neurons. Green fluorescence signals indicate neuronal activity. When an entire brain is captured at single-cell resolution at many volumes per second for minutes, neurons must be accurately segmented and individually tracked over thousands of image volumes. We sought an end-to-end analysis pipeline that would automate the extraction of neuronal activities from brain-wide imaging in *C. elegans*.

### Coarse alignment

The brain of a moving animal exhibits substantial rotations, translations, and deformations. The first step in our image analysis is the coarse global alignment of the 3D images at the scale of the whole brain. Accurate global alignment facilitates both manual annotation and proofreading by reducing differences in neuronal positions at different time points.

We trained a convolutional neural network (CNN) with the U-Net architecture [39] to perform a coarse global alignment of each recording. Once trained, the neural network works well on datasets from different animals (see Appendix I). This global alignment CNN (GA-CNN) automatically recognizes the points corresponding to the anterior, posterior, and central axis of the brain. Brain volumes can then be aligned across image volumes using point-set registration of the anterior, posterior, and axis coordinates. Global alignment is a common problem in image analysis, and alternative methods – such as identifying landmark neurons, multipole matching, *OpenCV* tracking [40] – would also work. At this step, we also reduce imaging noise by applying a Difference-of-Gaussian (DoG) filter to each image volume.

### Targeted augmentation

After coarse alignment and noise reduction, image volumes enter a pipeline that performs targeted augmentation of ground truth annotations for machine learning. This pipeline is illustrated in Fig. 2 A.

### Ground truth image selection and manual annotation

The first step is to select a small number of image volumes for ground truth (GT) manual annotation. These annotations will be used to train an initial CNN (iCNN) that segments and tracks neurons across the different postures that the brain can assume. At this step, it is useful to select image volumes corresponding to diverse postures, either individual volumes taken at regular intervals or a sequence of volumes when the animal exhibits substantial movement.

There are two useful ways of labeling neurons computationally for segmentation and tracking. One may label a neuron by a ‘key-point’, one chosen pixel inside the volume of the neuron. This is particularly convenient, for example, when only the nuclei of neurons are fluorescently labeled since neighboring nuclei are generally well separated and the correspondence between the key-point and the volume of the nucleus is easy to make. One may also label a neuron as a 3D volume, where all pixels in the volume of the neuron are used to identify the neuron. For simplicity, we will assume that we wish to perform key-point tracking in the following, and discuss the relatively small differences with 3D volume segmentation and tracking in the section ‘Segmenting and tracking volumetric objects’.

Identifying key-points can be partly automated, for example, by identifying local maxima of fluorescence in each image volume. When creating the GT manual annotation, non-rigid point-set registration can be used to track key-points across selected image volumes. Tracking by point-set registration is reliable when the worm is nearly immobilized but requires substantial manual correction when the worm is moving (Fig. 1).

**Figure 1.**
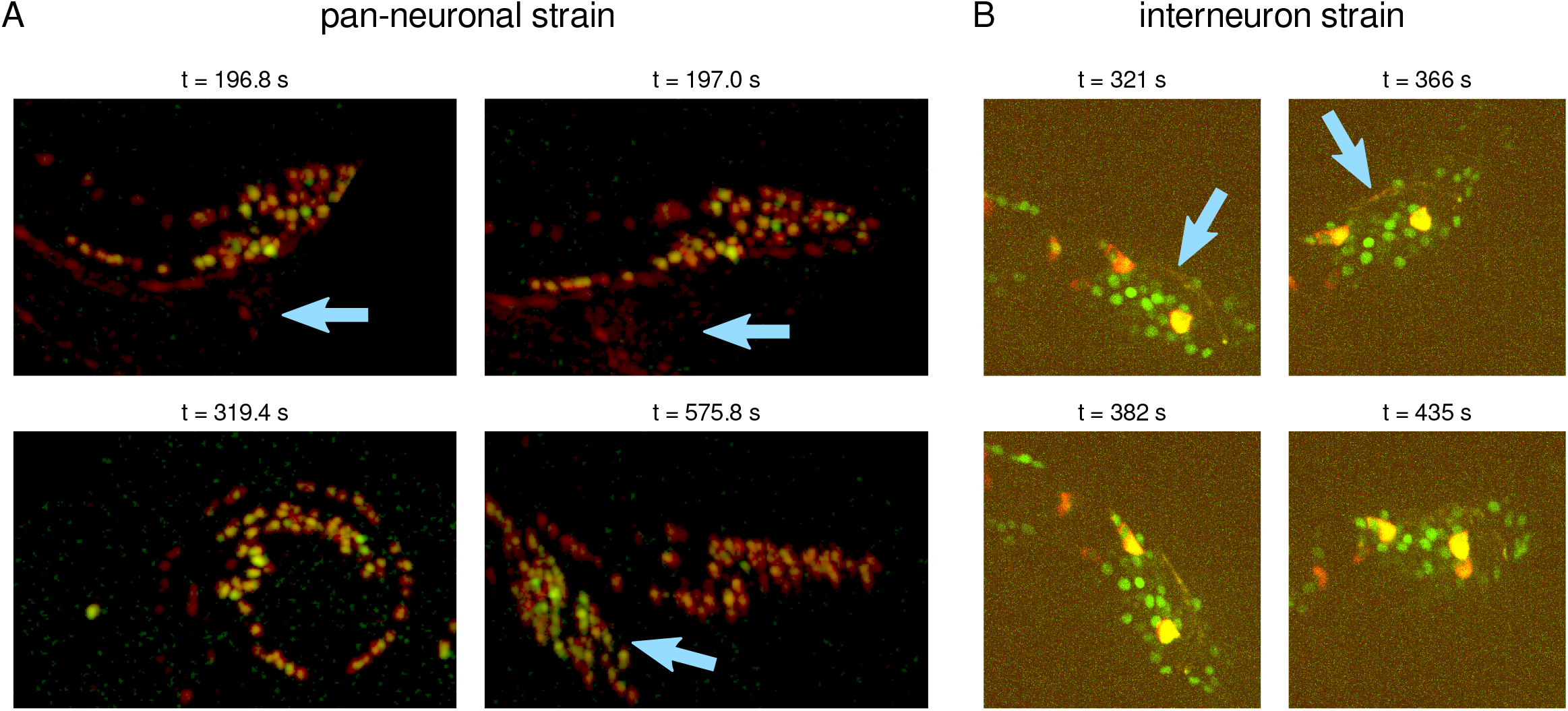
3D volumes from typical multi-neuron or whole-brain recordings of freely moving *C. elegans* worms after maximum intensity projection along the z-direction. A) The tail of a male worm with pan-neuronal nuclear GCaMP6s and red fluorescent proteins is shown in the presence of another, unlabeled hermaphrodite worm, indicated by arrows in the two top panels. In the bottom right panel, the head of the male worm, which is indicated by an arrow, appears in the field of view, next to the tail that is to be tracked. B) The interneuron strain carries a mixture of nuclear and cytosolic markers. The arrows indicate thin neurites.

After key-points for all neurons are selected across specific image volumes, they become the set of GT manual annotations.

### Training the iCNN

Using the GT manual annotations, we trained an initial CNN (iCNN) to automatically identify a small spherical region of interest around each key-point. We then used the iCNN to make a first set of predictions of key-point locations across image volumes.

We implemented the iCNN in a custom architecture that we call the 3D Compact Network (3DCN) (Figs. 2 B, S1). We designed the network to associate information over large distances when predicting neuron segmentations and key-point locations. We downsample after every two convolutions to associate information over large distances with a limited kernel size. This is similar to the downward branch of the U-Net [39]. To avoid increasing the raw size of the convolutional kernel and the corresponding increase in the number of fitting parameters for a bigger receptive field, we employ Atrous Spatial Pyramidal Pooling (ASPP) introduced in ref. [41].

The output of the iCNN should be a 3D image containing all segmented and tracked neurons with the same spatial resolution as the original image. A standard way to convert a low-resolution image in the latent space representation of a CNN back to its original resolution is to apply ‘upconvolutional’ layers with trainable weights. However, we found that simple tricubic interpolation of the latent space representation to estimate a high-resolution output was faster and as accurate as using up-convolutional layers. Our architecture thus resembles the FCN presented in ref. [42] for the upsampling process.

**Figure 2.**
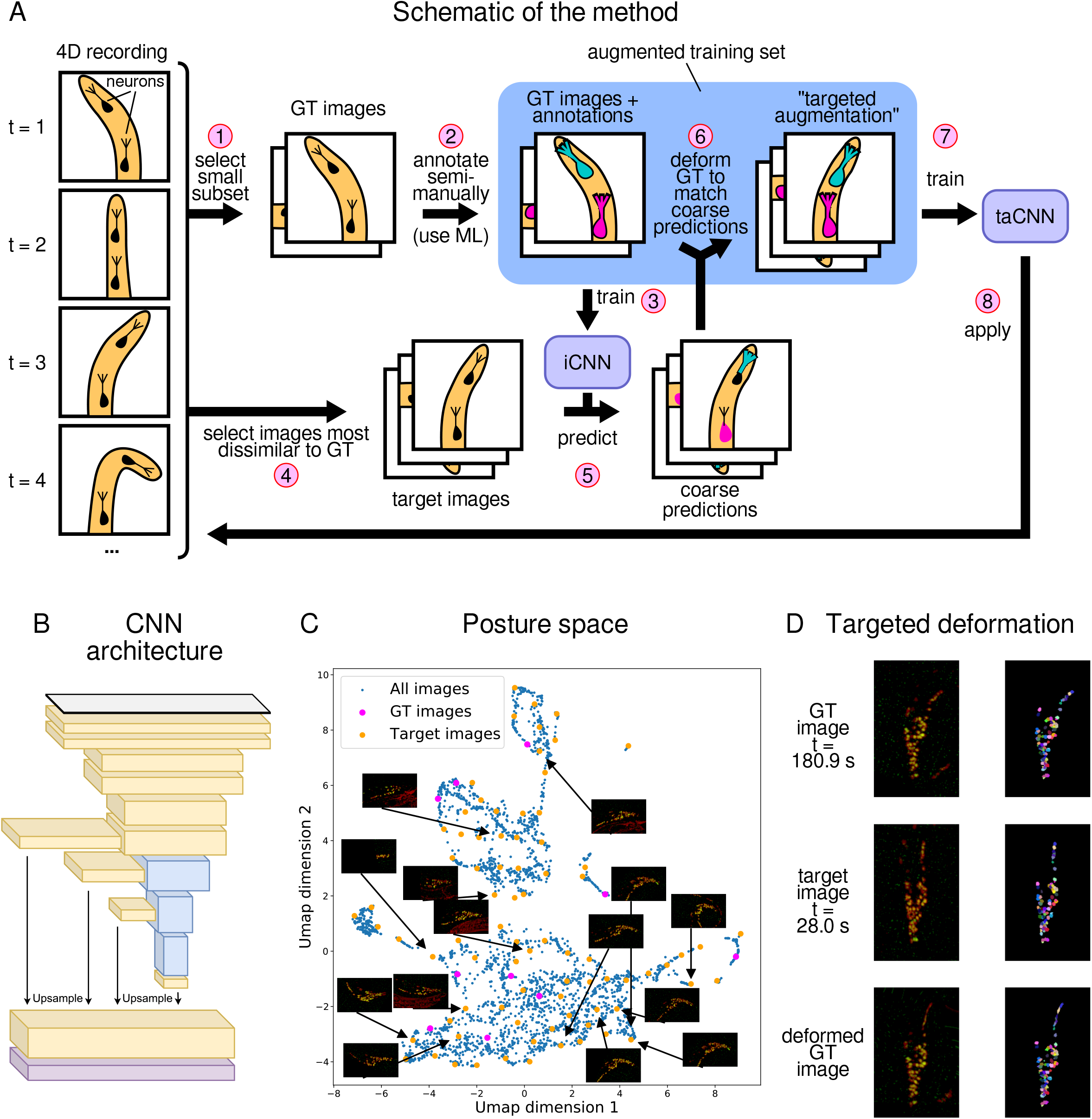
Illustration of the method. A: Steps involved in tracking. GT = ground truth, ML = machine learning. B: The CNN architecture used for the initial and augmented CNNs. C: The planar embedding of 3D images from the recording allows the similarity between 3D images to be measured by the Cartesian distances between their point representations in the plane. The embedding is performed by compressing the 3D image by an auto-encoder and mapping the latent space representations in the ‘bottleneck’ layer onto a plane using UMAP [36]. The representation of all (blue), GT (magenta), and target (orange) 3D images from a recording are shown. D: Example of the deformations performed on a GT image to match the target image. Left: maximum intensity projection of 3D images, right: initial CNN annotations of the images, which were used to perform the deformations

We allowed for the possibility of un-annotated neurons in each iCNN prediction because, when tracking the brain across behavior, some neurons can drop out of the field of view or be difficult to visualize at some time points. To allow the network to be able to ignore un-annotated neurons, we chose a cross entropy loss function which is able to mask out the channels corresponding to such neurons.

We implemented the iCNN with the properties described above in the 3D Compact Network (3DCN) architecture (Figs. 2 B, S1). The 3DCN exhibited improved accuracy, stability, and speed over U-Net for our tracking task (Table 1, Fig. 3 C). The 3DCN was efficiently trained on a desktop work-station.

**Table 1.** Benchmarks. All fractions are in percent (%). Tests were run on data that was hidden during method development. All runs were trained until saturation of the eIOU (see extended methods). All methods assume a training set size of 10. Each value represents the mean *±* standard deviation. The standard deviation around the mean describes the variability when we sample from the GT training sets of the three different recordings. The value for “Variance of FF” represents the variation when we sample from different GT training sets for the same recording. Top table: Comparison of the iCNN to the taCNN. In the “Masked” case, a second worm that was in the FOV was masked out. The “Strict” case requires the key-point to be the closest point found from the actual neuron. “Close” condition is a simple distance threshold of four pixels between the actual and the predicted key-point. Middle table: We compare our method to the CPD method (Supplemental note 1). Bottom table: We compare our network architecture to U-Net [39]. We found that the 3DCN is more than twice as fast as well as more accurate and stable.

|  | Masked |  |  |  | Not Masked |  |  |  |
| --- | --- | --- | --- | --- | --- | --- | --- | --- |
|  | Strict |  | Close |  | Strict |  | Close |  |
|  | iCNN | taCNN | iCNN | taCNN | iCNN | taCNN | iCNN | taCNN |
| Found Fraction (FF) | 68.0 ± 18.4 | **82.2 ± 10.2** | 76.2 ± 18.2 | **91.4 ± 8.4** | 71.5 ± 16.9 | **81.6 ± 8.3** | 80.3 ± 17.1 | **91.2 ± 5.2** |
| Variance of FF | 6.8 ± 7.0 | **3.9 ± 4.7** | 7.8 ± 8.3 | **4.4 ± 5.3** | 7.8 ± 8.3 | **1.8 ± 1.0** | 8.9 ± 9.9 | **1.9 ± 1.1** |
| Mistake Fraction | 13.6 ± 6.1 | **12.3 ± 5.7** | 5.4 ± 3.3 | **3.1 ± 2.5** | 14.6 ± 6.6 | **12.9 ± 5.4** | 5.8 ± 3.4 | **3.3 ± 1.9** |
| Miss Fraction | 18.4 ± 18.4 | **5.5 ± 7.9** | 18.4 ± 18.4 | **5.5 ± 7.9** | 13.9 ± 17.9 | **5.5 ± 4.2** | 13.9 ± 17.9 | **5.5 ± 4.2** |
| Extra Fraction | 7.4 ± 3.5 | 9.9 ± 5.0 | 7.4 ± 3.5 | 9.9 ± 5.0 | 10.3 ± 5.3 | 10.2 ± 4.8 | 10.3 ± 5.3 | 10.2 ± 4.8 |

|  | Method |  |  |  |
| --- | --- | --- | --- | --- |
|  | Strict |  | Close |  |
|  | CPD | 3DCN | CPD | 3DCN |
| Found Fraction (FF) | 25.3 ± 12.4 | **82.2 ± 10.2** | 28.0 ± 12.9 | **91.4 ± 8.4** |
| Variance of FF | **1.7 ± 0.5** | 3.9 ± 4.7 | **1.8 ± 0.5** | 4.4 ± 5.3 |
| Mistake Fraction | 34.5 ± 4.0 | **12.3 ± 5.7** | 31.8 ± 4.6 | **3.1 ± 2.5** |
| Miss Fraction | 40.2 ± 8.9 | **5.5 ± 7.9** | 40.2 ± 8.9 | **5.5 ± 7.9** |
| Extra Fraction | 3.7 ± 1.8 | 9.9 ± 5.0 | 3.7 ± 1.8 | 9.9 ± 5.0 |

**Table 1.** Benchmarks. All fractions are in percent (%).
|  | Architecture |  |  |  |
| --- | --- | --- | --- | --- |
|  | Strict |  | Close |  |
|  | U-Net | 3DCN | U-Net | 3DCN |
| Found Fraction (FF) | 48.7 ± 8.6 | **76.2 ± 2.0** | 56.4 ± 9.8 | **87.2 ± 1.9** |
| Variance of FF | 8.6 | **2.0** | 9.8 | **1.9** |
| Mistake Fraction | 38.6 ± 6.4 | **14.7 ± 0.9** | 30.9 ± 7.6 | **3.7 ± 0.8** |
| Miss Fraction | 12.7 ± 4.5 | **9.1 ± 2.0** | 12.7 ± 4.5 | **9.1 ± 2.0** |
| Extra Fraction | 19.6 ± 1.1 | 16.1 ± 1.0 | 19.6 ± 1.1 | 16.1 ± 1.0 |
| Gradient Step Time | 0.486 ± 0.001 | **0.237 ± 0.004** | 0.486 ± 0.001 | **0.237 ± 0.004** |

**Figure 3.**
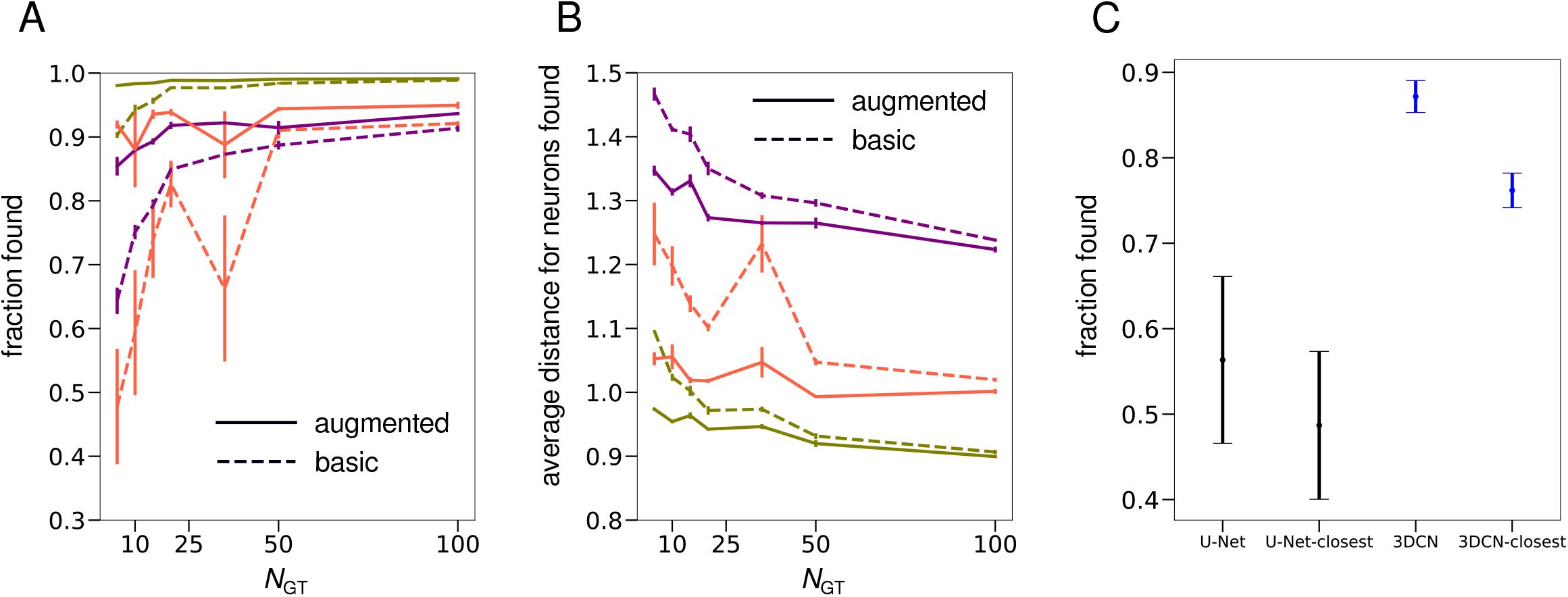
Evaluation of the performance of our pipeline for key-point tracking applied to three different recordings of 5-10 min duration (1500-3000 image volumes), each indicated in a different color (gold, orange, purple). A: A neuron was considered found if the predicted key-point was within 4 pixels of the ground-truth key-point. Solid lines indicate the performance of the augmented CNN, dashed lines indicate the performance of the initial CNN. B: Mean distance from the ground-truth for found neurons. Each recording is represented by the same color as in panel A. C: Comparison of 3DCN and U-Net. The ‘closest’ condition requires that the predicted key-point marking a neuron is the closest predicted key-point to the actual key-point.

### Selecting images for targeted augmentation

We wanted to minimize the number of manually annotated GT volumes that the pipeline would need. Our strategy was to enlarge and diversify the set of GT annotations by automatically generating synthetic GT annotations.

For this, the algorithm selects *target images*, an optimally representative subset of all images from the recording, which we used as templates for synthetic GT annotations. The images were selected to be different from the set of GT manual annotations and different from each other. To accomplish this, we needed a distance metric to estimate image similarity. We used a convolutional autoencoder [43] to create a low-dimensional latent space representation of all recorded image volumes. We reduced the autoencoder’s latent space representation further to two dimensions using the UMAP [36] method. The distance between two points in the UMAP plane is a measure of the overall similarity between the corresponding brain images (Fig. 2 C). In this way, the algorithm selects a set of target images that broadly samples the points in the latent space representation (See Extended Methods).

### Creating synthetic GT annotations

One way to create additional GT annotations for the selected target images would be to use the iCNN to make coarse predictions for neurons in the target images, and then perform proofreading and manual correction. Instead, we aimed for a less laborious and fully automated method by leveraging the GT images and their annotations. For each target image, the method selects in turn the most similar GT image, and the iCNN makes coarse predictions for neurons in the target image. Next, the goal is to deform the GT image to resemble the target image. For key-point annotations, we fit a low-frequency deformation field that optimally maps the key-points in the manually annotated GT image onto the coarse predictions of key-point locations in the target image (Fig. 2 D). (See ‘Segmenting and tracking volumetric objects’ for the modifications of this step for 3D volume tracking.) We implement this fitting by minimizing the mean L1 displacement of key-points after deformation and by restricting the Fourier modes of the deformation field to low frequencies. We used the L1 loss because it is minimally sensitive to the outlying errors made by the iCNN – the inaccurate coarse predictions of the iCNN that assign key-points far from their actual locations.

Thus our deformation field, 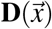, is

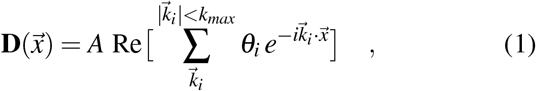

where 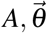 are the free fitting parameters. The loss function is

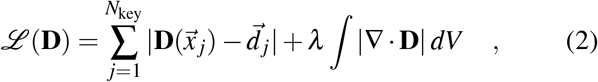

where *N*_key_ is the number of key–points and 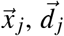 are, respectively, the ground truth location of the *j*^th^ key-point and the displacement vector from the latter to the corresponding ‘coarsely’ predicted location. The second factor is a regularization term to keep the divergence of the deformation field reasonable.

Finally, we used the fitted deformation field to generate both a synthetic image and its corresponding annotation by applying the field to both the GT image and its manual annotation. The synthetic image that results from this deformation will resemble the target image while propagating the correct annotations from the GT image. The resulting key-point locations represent a reliable annotation of the synthetic image even when the original coarse prediction made by the iCNN is unreliable, albeit with less resemblance to the target image. This synthetic image and its key-point annotations can then be added to the GT images and their manual annotations to create a larger training set for a ‘targeted augmentation CNN’ (taCNN) that has better performance than the iCNN because it covers more postures. The targeted augmentation CNN is then applied to all images in the recording.

### Evaluation of the targeted augmentation method

We evaluated our method for targeted augmentation by segmenting and tracking key-points in brain-wide recordings of the male brain in freely behaving *C. elegans* performing their mating ritual [12] (Fig. 1 A). We obtained four recordings, each containing 1500-3000 image volumes (5-10 min per recording). In the previous study of male mating behavior, all recordings were fully manually annotated, which required 100-200 hours per recording. We asked whether our technique using the taCNN would achieve comparable reliability as full manual annotation, but with a smaller set of images that needed to be manually annotated to train the neural network. We developed our technique using one brain-wide recording of the male brain. We evaluated the performance of our method by comparing the results of taCNN predictions to the results of full manual annotation of the three other ‘held-out’ recordings (Table 1, Fig. 3).

In practice, we found that we could set the number of target images from a brain-wide recording to *N*_target_=80, which saturated the accuracy of the final taCNN predictions in tests of our method (Fig. S2). The utility of our technique is measured by how much it reduces the number of manually annotated GT images that are needed to achieve the same accuracy in segmenting neurons and tracking key-point locations as the iCNN. We found that the taCNN is able to identify nearly the same fraction of neurons across the three held-out brain recordings when trained with only 5-25 manually annotated GT images as the iCNN that is trained with 100 manually annotated GT images (Fig. 3 A). This did not change when we varied the pixel-distance for the threshold determining whether a key-point was called correctly; the taCNN always substantially outperformed the iCNN (Fig. S3). Furthermore, when neurons where found, they were identified more accurately with targeted augmentation (Fig. 3 B).

In image volumes containing the male brain during mating behavior, the hermaphrodite often entered the field of view, a decoy that challenged both the iCNN and taCNN. The taCNN out-performed the iCNN in segmenting and tracking neurons in the male brain whether or not we erased the decoy hermaphrodite from image volumes. If we increased the threshold for a correctly identified neuron – by requiring not only that it has to be within a given pixel-distance of the actual neuron but also that it is the closest predicted neuron – the taCNN still always outperformed the iCNN (Fig. S4).

We compared segmentation and tracking by taCNN with another commonly applied method, Coherent Point Drift (CPD) point-set registration [44] (Supplemental note 1). Point-set registration requires fully segmented neurons in all image volumes, whereas the taCNN performs both segmentation and tracking. CPD tracks neurons by finding the optimal correspondence between neuron locations between two images. We applied CPD to our brain-wide imaging datasets of male mating behavior. We created reference sets of different sizes so that CPD could match neurons between a new image and the closest image in the reference set. The larger the reference set, the better CPD should work, analogous to increasing the size of the training set for the taCNN. We found that CPD accurately tracked fewer than 20% of neurons for similar reference set sizes where taCNN correctly tracked >80% of neurons (Fig. S5).

### Segmenting and tracking volumetric objects

We asked whether the taCNN could be used to segment and track the 3D shapes of neurons, not just key-point locations. In *C. elegans* and other animals, calcium dynamics is often different in different parts of a cell in functionally important ways (e.g., soma vs. neurites). [26] These dynamics are missed when recording calcium dynamics with nuclear-localized probes, and require reconstruction of the spatial distribution of calcium dynamics in different neuronal compartments.

We developed a transgenic strain to measure calcium dynamics throughout cytosolic compartments in interneurons of the hermaphrodite nerve ring that are responsible for chemosensory processing, including the AIY interneuron and RIA interneuron that primarily exhibit calcium dynamics in their neurites and not their cell bodies. This strain (SJR15) expressed red fluorescent proteins (wrmScarlet and mNeptune) and GCaMP6s in the cytosols of the AIA, AIY, AIZ, and RIA interneurons. This strain also expressed red fluorescent proteins in the nuclei of the AIB, RIB, and RIM interneurons, and GCaMP6s in the nuclei of all neurons in the nervous system.

We modified some steps of the taCNN method to segment and track the shapes of neurons and nerve fibers in image volumes. First, we needed to generate GT annotations of neuronal shapes and structures. We began by using adaptive thresholding to identify contiguous fluorescently-labeled objects within each image volume. All objects are associated with a number of quantitative measures – e.g., overall size, aspect ratio, brightness – that can be used as identifiers. We quantified a large set of these geometric features for all objects across all image volumes. We performed k-means clustering on the set of geometric features, resulting in the automated clustering of the same objects found in different image volumes. In effect, we used k-means clustering as an elementary method for tracking objects that had reasonable accuracy (approx. 60%, see Methods). Finally, the volumetric structures that were automatically segmented by adaptive thresholding and tracked by k-means clustering were then manually proofread and corrected, a step that was much faster than their full manual annotation. Thus, for volumetric object tracking, we used machine learning already at the manual annotation step.

As for key-point tracking with the taCNN, we augmented the training dataset for neuronal shapes by adding synthetic GT annotations for a set of target images that were created by deforming similar manually annotated GT images. Instead of fitting a low-frequency deformation field, we obtained better results by applying a non-linear optical flow transformation [45, 46] from each GT image to each target image.

Performance evaluations of the taCNN applied to volumetric tracking of neuronal shapes and structures agree with evaluations of key-point segmentation and tracking (Fig. 4). Using the taCNN led to substantial improvements in comparison to the iCNN in the fraction of neurons found (Fig. 4 A). We further compared the accuracy in the segmentation of individual neuronal objects by computing the intersection-over-union (IoU) of all objects found. By this measure, the iCNN and taCNN performed similarly, with the taCNN slightly outperforming the iCNN for small training sets and vice versa for medium and larger training sets (Fig. 4 B, note y-axis units). As with key-point tracking, using the taCNN substantially reduced the amount of manual annotation and proofreading. With targeted augmentation, small training sets (*N*_GT_ = 5) produced results similar to training sets that were 3-5 times larger and that were not enhanced by targeted augmentation.

**Figure 4.**
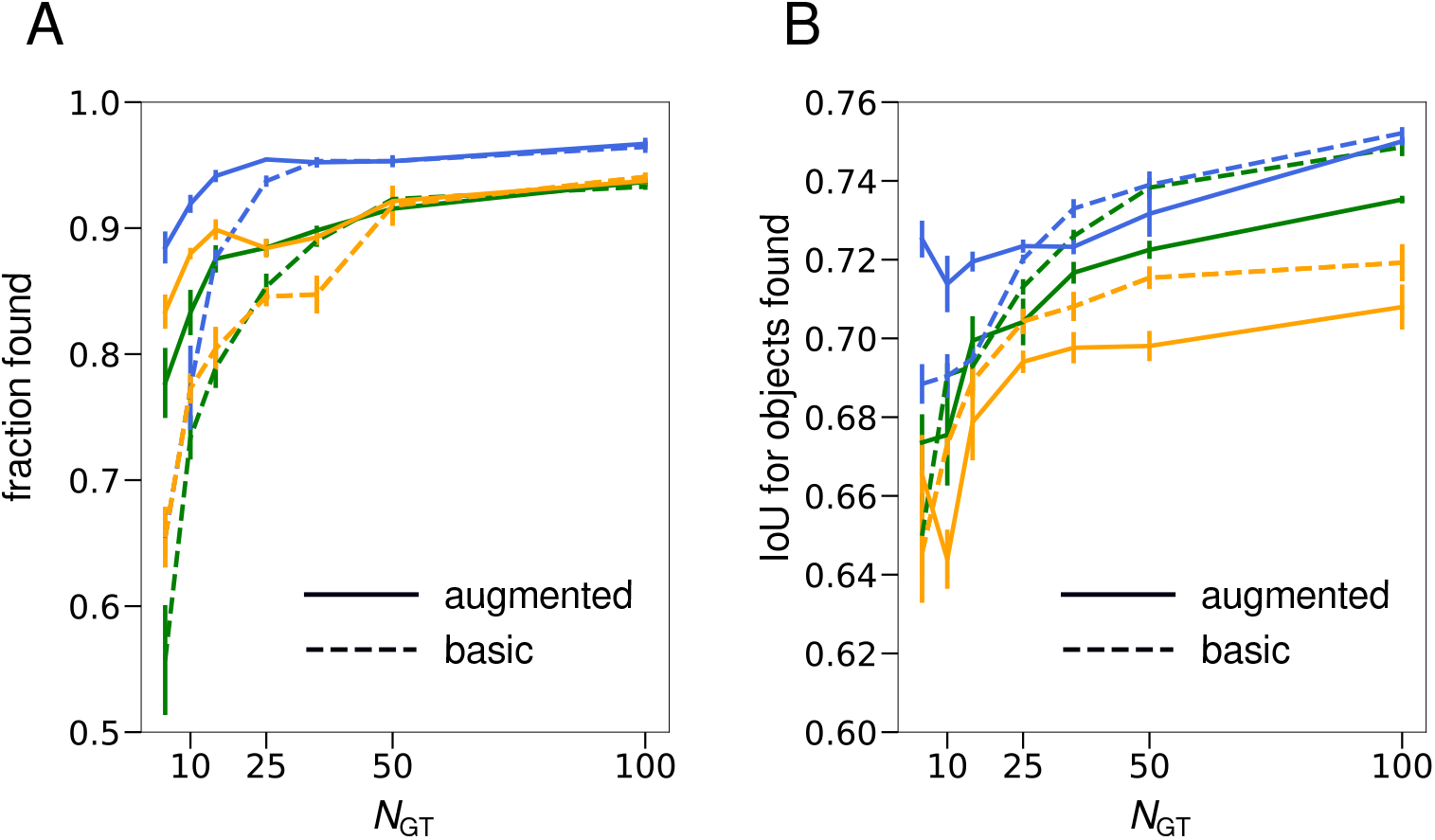
Evaluation of the performance of our pipeline for 3D volume tracking, applied to three different recordings of 10 min duration (1715 image volumes), each represented by a different color (blue, green, yellow). A: For 3D volumes, we considered objects as found if the IOU between the predicted and the manual annotation exceeded 0.33, i.e., the intersection of two 3D objects was greater than the volume of the ground-truth or the predicted 3D volumes. Solid lines indicate the performance of the augmented CNN, dashed lines indicate the performance of the initial CNN. B: Intersection-over-Union (IoU) for 3D volumes. Each recording is represented by the same color as in panel A.

### Coupling of sensory information to interneuron activity

To apply our method to measurements that could not be easily analyzed with previous methods, we recorded calcium activity in the second-layer interneurons RIA, RIB, and RIM. Worms expressed nuclear localized GCaMP6s pan-neuronally (Fig. 5 A). Additionally, RIB expressed nuclear-localized mNeptune, RIM expressed nuclear-localized wrmScarlet, and RIA expressed cytosolic GCaMP6s and mNeptune. The different choices of red fluorescent protein allowed us to distinguish the nuclei of RIB and RIM neurons. Worms were placed in microfluidic chips encompassing a structured arena, adapted from ref. [47], and were pulsed with IAA or 2-nonanone medium pulses of 20 sec duration and 1 min period. The worms were imaged as described above, and activity was extracted after analysis with our pipeline.

**Figure 5.**
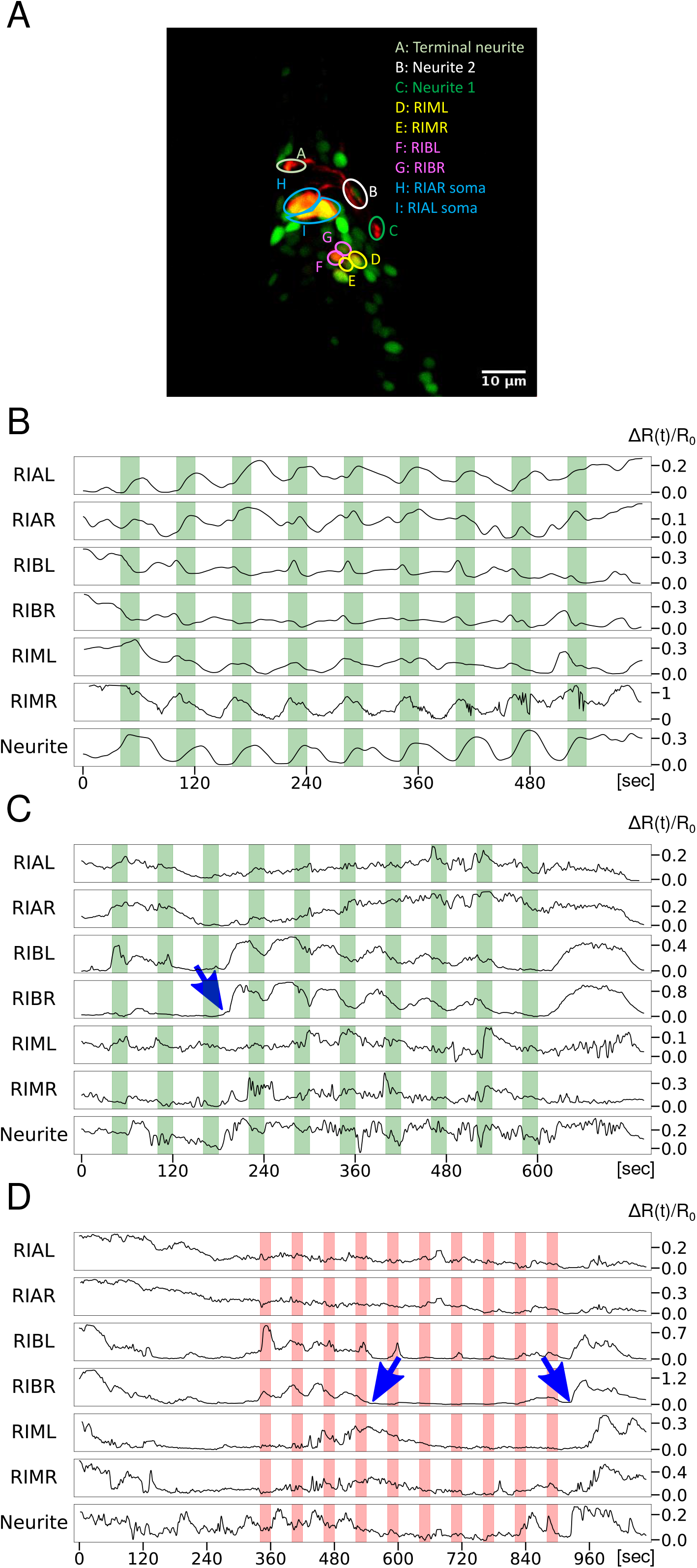
Calcium activity in the second-layer interneurons RIA, RIB, and RIM. “Neurite” indicates a segment of the neurite of RIA. A: Maximum intensity projection of a 3D image of a worm expressing nuclear-localized GCaMP6s pan-neuronally, nuclear-localized mNeptune in RIB, nuclear-localized wrmScarlet in RIM, and cytosolic GCaMP6s and mNeptune in RIA. B-D: Traces of GCaMP fluorescence divided by red fluorescence (*R*(*t*)), from which the baseline *R*_0_ was subtracted and the difference was normalized by *R*_0_. Green bars indicate 2-nonanone pulses. Red bars indicated IAA pulses. B: Entrained calcium activity in all three neurons. C: Entrainment begins after the third odor pulses, indicated by the arrow. D: Arrows indicate when entrained activity stops and when activity restarts later.

Periodic stimuli are powerful tools for elucidating circuit behavior. [33, 34] We observed a number of rich activity patterns in freely behaving *C. elegans*. Neuronal activity in the second-layer interneurons, which are thought to be closely linked to locomotion, could be entrained by the odor pulses (Fig. 5 B). However, this varied not only from worm to worm but also from time to time for the same worm. Some animals showed full entrainment to the external odor pulses while others exhibited none. Interestingly, worms could switch entrained activity on or off (Fig. 5 C, D). These observations suggest that long recordings of single animal activity in multiple neurons continue to reveal novel phenomena, highlighting the importance of efficient 3D image analysis techniques.

### Graphical user interface (GUI)

We created a python-based GUI for viewing and annotating 4D recordings, launching the steps of the pipeline, including targeted augmentation and the neural network training, and for viewing and proofreading the results of the predictions of the different neural networks, iCNN and taCNN (Fig. 6). The user can leaf through z-stacks, view z-projections, and annotate by placing or moving key-points as well as drawing 3D masks with a cubic pencil or by local thresholding (‘3D bucket fill’). A ‘neuron bar’ and a dashboard are designed to make it easier for the user to spot incomplete annotations and navigate long recordings.

**Figure 6.**
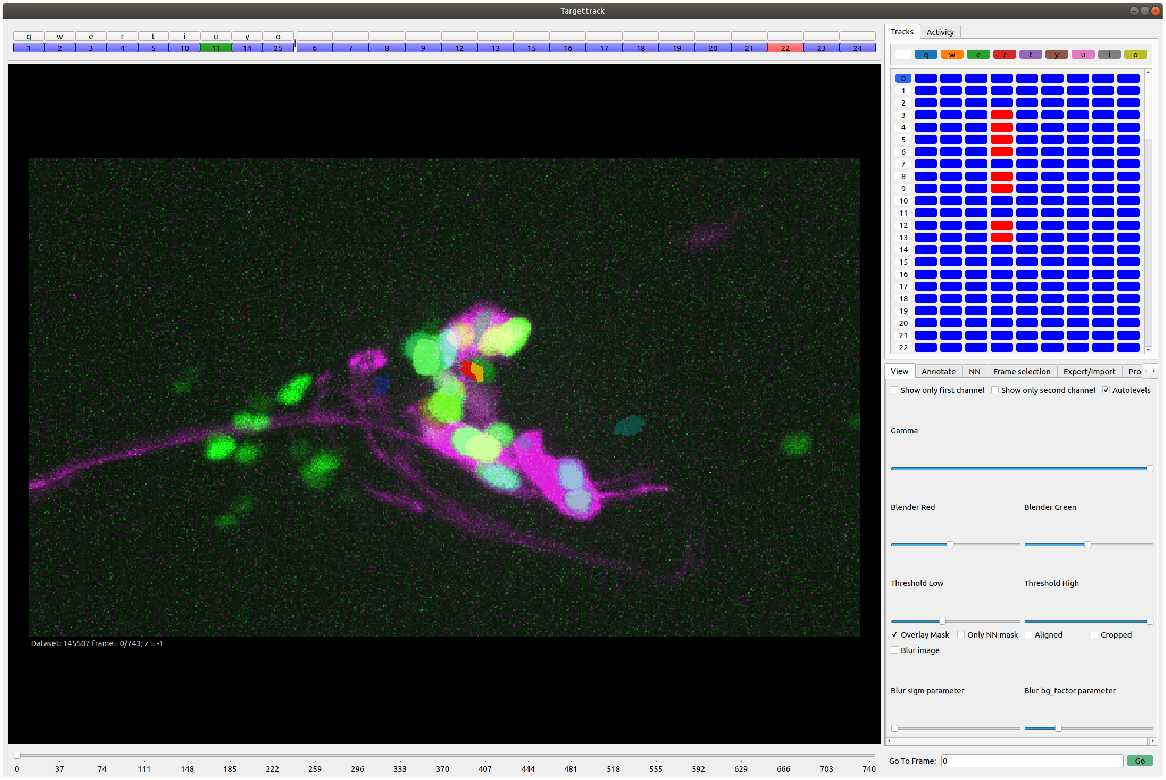
Targettrack GUI for viewing and annotating 4D recordings as well as applying the pipeline.

## Discussion

Many laboratories now perform whole-brain or multi-neuron imaging with single-cell or subcellular resolution in different animals [5, 12, 37, 38]. The current bottleneck is data analysis: converting large-scale recordings of image volumes over time into the segmentation and tracking of individual neuronal activities throughout the brain. Manual annotation is the most reliable way of analyzing brain-wide recordings. However, manual annotation becomes unacceptably labor-intensive when recordings become numerous, long, or encompass many neurons. Manually annotating even a few minutes of recording can take hundreds of hours [12], slowing progress.

Machine learning is ideally suited to the pattern recognition task of segmenting and tracking neurons. However, any method must deal with substantial image-to-image variability due to optical changes, biological differences, and movement and deformation during animal behavior in multi-neuron recordings. To be accurate, neural networks must be trained with representative images and annotations that span the range of image variability [21]. As the diversity of images and the number of neurons increase, the amount of training data that is required also increases, which represents a significant burden if training data is produced by manual annotation. After automated segmentation and tracking using a neural network, manual proofreading also becomes a burden if the network has high error rates. Thus, traditional machine learning techniques involve a trade-off between the amount of manually annotated data that is used to train a neural network and the amount of manual proofreading needed to correct errors.

We present innovations that allow a CNN to both minimize the required amounts of manually annotated training data and of proofreading. We optimized the neural network architecture to reliably segment and track neurons within a rapidly moving and deforming *C. elegans* brain. Our method generates part of its own training data based on a small number of manually annotated images using targeted augmentation. By estimating the deformation of a brain volume in a target image based on a manually annotated image, we automatically create new synthetic images with reliable annotations. When a CNN is trained with a small number of manually annotated images along with diverse synthetic images and annotations, its reliability increases substantially, reducing the amount of proofreading needed.

The automatic generation of synthetic training images has previously been explored in the medical domain. For example, [48] focuses on segmenting cardiac, prostate, and pancreas images, and generates synthetic training examples using GANs. In contrast, [49] focuses on the segmentation and registration of brain and knee MRIs. The method uses a neural network to learn displacement fields between images. The closest approach to ours may be that of ref. [50]. This method segments brain MRIs with the help of synthetic training data generated by fitting free-form deformations between existing labeled and unlabeled images. Overall, the task in the present work, i.e., tracking freely moving worms, is quite different from segmenting MRI images, among other reasons, because the 3D volumes within a recording are much more heterogeneous. This makes segmentation and tracking of our recordings substantially more challenging.

Because of experiment-to-experiment variability, any image analysis method will be more reliable when there is ground-truth training data specific to each experiment. Our pipeline starts with a small number of manually annotated GT images for each experiment and requires no extraneous information. The pipeline effectively learns the brain-wide deformations that occur in one individual experiment. It then automatically expands the originally annotated GT images into a larger training dataset using the learned deformations. Our method is ideally suited for use by different laboratories and changing experiments, as it flexibly adapts to the specific imaging conditions of each individual experiment.

Another noteworthy aspect of our targeted augmentation pipeline is that it is germane to 3D images. While 3D images may be perceived as merely more difficult to analyze than 2D images because of their larger sizes, they also afford additional opportunities for image analysis. In three dimensions, the worm brain can in principle be mapped by deformation from one image to any other. This is not true for 2D projections of the brain, where, for example, two crossing lines cannot be uncrossed by deformation. Thus, there may be additional unexplored opportunities for simplifying image processing tasks in 3D.

We have developed and applied our approach to particularly challenging problems in *C. elegans* brain-wide imaging. For example, during its unrestrained mating behavior, the posterior brain of the male nematode exhibits rapid and dramatic movements and deformations as the male interacts with a hermaphrodite, itself a visual object that distracts and challenges the performance of the neural network that is focused on the male. In practice, we have required 200 hours to fully manually segment and track 76 neurons in the male tail in just one 10 minute recording. Using our taCNN pipeline from end-to-end on the same dataset, we reduced the amount of manual effort to 65 hours – 5 hours to generate a small but adequate amount of manually annotated GT images and 60 hours to comprehensively proofread all automatically segmented and tracked images. The latter is generally difficult to reduce as proofreading is still necessary even for simpler image analysis problems with even lower error rates [51]. Thus, the speed-up of our method expands what is feasible for brain-wide imaging of small-animal models in neuroscience.

We expect the effectiveness of our pipeline to improve with advances in imaging. Segmenting and tracking neurons will become easier with better image quality and higher spatial and temporal resolution, which will be possible with improvements in microscopy and fluorophores. Multi-color imaging approaches will allow the taCNN to incorporate more information that will facilitate its reliability. In this work, we have not used any information beyond a single fluorescent channel to keep the method as general as possible. One strength of the CNN framework is that it is straightforward to add additional types of information such as additional image channels.

## Methods

### Initial coarse alignment of whole-brain images

To facilitate later steps in the segmentation and tracking pipeline, the algorithm performs an initial coarse alignment of all whole-brain images in each recording. There exist many techniques for the coarse alignment of images, two of which we present here.

#### Whole-brain recordings of neurons segmented and tracked as key-points

When tracking many neurons inside freely moving *C. elegans*, our tests were applied to a set of manually annotated recordings of the posterior nervous system of the male during mating behavior [12]. We used these manual annotations to train a 2D U-Net to solve the problem of coarse alignment of whole-brain images. To perform coarse alignment, an effective algorithm must *(a)* determine which pixels in a 3D image correspond to the brain and *(b)* determine the brain’s orientation. We created a training dataset for the 2D U-Net by converting the comprehensive manual annotations of segmented and tracked neurons from previous work [12] into a simple map of neuron locations within the brain distinguished by their coordinates along the anterior-posterior axis. After training, the 2D U-Net was able to identify neurons and estimate their coordinates when given a new 3D brain image. Next, the algorithm computes the gradient of these estimated coordinates, which represents the orientation of the worm brain in each new image. Finally, this computed orientation is used to perform an affine alignment of each image. We found that the 2D U-Net network, when trained with recordings of 1-3 animals, was effective for identifying neurons in images of other animals and thus is useful as a general tool for coarse alignment.

#### Recordings of neurons segmented and tracked as 3D volumes

We used a different coarse alignment procedure to orient neurons represented as 3D shapes. The algorithm identifies a few landmark neurons, that is, particularly bright neurons that are visible in all 3D brain images and that can be automatically detected using a high threshold on the fluorescence intensity. We then compute the coarse alignment by performing the non-rigid Jian-Vemuri [52] point cloud registration algorithm. We approximate the point-wise registration of landmarks by rotations and translations of the images in the x-y plane.

### Ground-truth image selection

To train the 3DCN to perform segmentation and tracking of neurons, we need a diverse set of manually annotated ground truth (GT) images. The GT images need to be diverse to account for the different postures of moving animals across a particular recording. We either select images at regular intervals throughout the recording or select an easy-to-annotate image sequence where the animal moves substantially. Either method was sufficient, and thus we did not develop a computational method to select GT images. In practice, the user may benefit from flexibility when choosing images for GT manual annotation. An algorithm that prescribes the 3D images to annotate may not, for example, be able to account for the varying subjective difficulty of annotating particular 3D images.

### Ground-truth annotations

Once GT images are selected, they need to be accurately annotated to serve as training datasets for the 3DCN. For freely moving animals that are to be tracked by key-points, we did not automate the annotation of GT images. For example, non-rigid point-set registration methods such as CPD [44] (Supplemental note 1) were not sufficiently accurate to speed up manual annotation. Thus, all GT images for segmenting and tracking neurons in freely moving animals are obtained by manual annotation. For semi-immobilized worms, however, CPD pointset registration can generate rough annotations that can be proofread in less time than full manual annotation.

For segmenting and tracking the 3D shapes of neurons, we developed a semi-automated method for the annotation of GT images. First, we segment the images, that is, identify the 3D structures representing the nuclei, soma, and neurites of fluorescent neurons. In each recording, we enhance each 3D image by applying a Difference-of-Gaussians filter. We apply a threshold to the enhanced image to keep a percentage of the brightest pixels. We compute the Euclidean distance transform of the thresholded image, which is then smoothed with another Gaussian kernel. The local maxima of the smoothed distance transform are used as seeds for a watershed algorithm, which finds the connected volumes around each local maximum. We applied a user-defined threshold to discard small local maxima that are too close to bigger maxima. We merge volumes that were overlapping or adjoining. We adjust the algorithm to remove segmentation errors (by merging pieces of the same object that were erroneously split into different volumes) without creating merge errors (avoiding the erroneous merging of different adjoining objects). To do this, we only merge when the contact areas are large, that is, when the overlapping/neighbouring surface divided by the smaller volume (to the power of 2/3) is greater than a user-defined threshold. Volumes that are too small are excluded.

In addition to segmentation, we created complementary tools for the manual creation and deletion of 3D annotations in the GUI. This tool allows 3D volume masks to be drawn with a cubic ‘pencil’ or created by local thresholding.

Once all 3D individual objects are segmented and identified by adaptive segmentation and manual annotation, each is represented by a vector of quantitative features. These features are volume, total fluorescence intensity, maximum intensity, variance of intensity, ratio of diameter to volume, and eigenvalues of the moments of inertia matrix. We then apply K-means clustering to segregate and locate all 3D objects in the feature space. When applied to different time points and different 3D images, K-means clustering should assign the same objects to the same locations in the feature space. We found that the accuracy of this method was about 60% (the average number of correctly tracked objects in 12 frames in one recording). Proofreading and correcting the results of this elementary tracking yielded the GT annotations.

### Point-to-mask conversion

For key-point tracking, the method should predict a key-point corresponding to the location of each tracked neuron. During the training of the neural network, however, it did not suffice to supply a single pixel to be predicted for each neuron. Instead, for each annotation, we generate a mask in which all pixels within a radius of 4 pixels from the ground-truth keypoint are labeled as the neuron which is to be predicted. So, the neural network is trained to predict a 4-pixel ball of pixels to identify a neuron. When the neural network is then applied to new images in the recording, we straightforwardly reduce the set of predicted label pixels to a single key-point pixel (see ‘Post-processing’ below).

### initial CNN

Once we have an initial set of GT images, we use them to train an initial CNN. The architecture of the CNN, which we call 3DCN, is illustrated in Figs. 2 C, S1. Several features make our CNN more accurate, as well as faster to train and apply than the popular U-Net [39].

First, we accounted for anisotropic resolution. In most 3D light microscopy – whether confocal, two-photon, or light-sheet microscopy – the resolution in the xy directions is higher than the resolution in the z-direction. We thus applied 2×2×1 down-sampling to compensate for the difference in xy- and z-resolution.

We found that the 3DCN did not need to train the upsampling layer to generate the final predictions. Instead, we found that a simple tricubic interpolation was an effective and computationally efficient way to extract predictions.

We found that the trained network needed to account for long-range correlations to accurately identify neurons. Deformations in distant parts of the animal contain information about the animal’s overall posture. To capture long-range information without large kernel sizes, we employed atrous convolutions in the ASPP module [41]. In brief, this method enlarges the field of view of the kernel by skipping over features that are adjacent to features already captured.

### Target set for augmentation

For targeted augmentation, the algorithm selects a set of diverse 3D images from the recording as templates for deforming the GT images and annotations. This step requires image volumes to be compared and their similarities to be quantified. To compute the similarity between any two images, all 3D images from each recording are used to train a convolutional autoencoder. Thus, the autoencoder maps each 3D image to a compressed representation in the network’s latent space. In principle, the relevant information from the 3D images, e.g., noise, intensity changes, worm body deformations, other objects in the field of view, and the field of view, are captured in the latent space. We used the L2 loss function so that the network focused on the bright part of the image rather than trying to summarize the background noise distribution, which is a majority of the pixels. Because most deformations are in the x-y plane, it sufficed to perform a maximum intensity projection in the z-direction first, and apply a 2D autoencoder. After normalizing the latent vectors from the autoencoder to zero-mean and unity-standard deviation, the representation of each image was mapped onto a plane using UMAP [36].

### Targeted augmentation

Targeted augmentation succeeds when the coarse annotations by the initial CNN, while not accurate enough to be satisfactory as the final results of the pipeline, suffice to deform the GT annotations to match the target images. We use the coarse annotations to match the nearest GT image to each target image by computing the most effective and smoothed deformation field. For key-points, we compute the deformation field with a wavelength cut-off in Fourier space by fitting the vectors pointing from the neurons in the GT annotation to the neurons predicted in the target image by the initial CNN.

For 3D objects, we compute the deformation field in multiple steps. First, we perform a better coarse alignment of the GT image and the target image. To do this, all objects in the GT image and all predicted objects in the target image are approximated as clouds of points. After this, all objects are matched with the Jian-Vemuri [52] point cloud registration method. The Jian-Vemuri method [52] ignores the identities of the neurons and matches the constellation of point clouds. It generates vectors representing the match of each point in the point cloud in the GT image to a corresponding point in the point cloud in the target image. Our algorithm then approximates the transformation represented by these vectors as an overall translation and rotation of the whole GT image. Subsequently, optical flow [45, 46] finely and non-rigidly registers the rotated and translated GT with the target image. The deformation generates a new GT image and annotation which are often close to a valid annotation of the target image. The deformation field, which represents the translation-rotation and the optical flow registration, can then also be applied to the annotations in the GT image. To prevent objects in the annotation from being torn due to the optical flow step, we post-process them by computing the nearest *α*-shape (based on a radius of 5 pixels) that corresponds to each individual object in each 2D mask slice [53].

To explain the improvement of the annotations by the augmented CNN compared to the initial CNN, we speculate that even when the deformations are small, the deformed images force the neural network to have a more consistent representation of the input images. If the results of the deformation are imperfect, the deformed image and annotations are not expected, in principle, to harm the neural network performance. When this happens, the augmentation just fails to increase the diversity of realistic 3D brain postures. The validity of this idea can be quantitatively assessed based on the data presented in Figs. 3, 4.

### Post-processing (not shown in Fig. 2)

For key-point tracking, the neural network predicts a set of points as labeling a neuron, not a single key-point pixel. This is because the training is performed with a 4-pixel ball around each key-point annotation (see ‘Point-to-mask conversion’). Consequently, to generate key-points from the predicted labels, we take the largest connected components of predicted pixels for each neuron, and calculate its ‘center of mass’ where the weight of each pixel is given by the fluorescence intensity in the recorded image. The center of mass is then assigned as the predicted key-point. This step also addresses the problem that sometimes the neural network predicts disconnected pixels to label an individual neuron; by taking the largest connected component, stray pixel labels are ignored.

For 3D volumes, the neural network sometimes mislabels individual pixels that are part of one neuron as belonging to another neuron. Thus, we check all pixels that are part of the same connected component, and if there are pixels of one neuron touching or inside another neuron, they are merged with the larger object.

### Strains

For imaging interneurons, we used ADS1001 [P*rgf-1:NLS-GCaMP6s*] from ref. [12], which expresses nuclear-localized GCaMP6s pan-neuronally. For recordings of second-layer interneurons, *sjxIs9* was generated by integrating *sjxEx9*[P*glr-3::mNeptune::GCaMP6s*; P*sto-3::NLS-mNeptune*; P*cex-1::NLS-wrmScarlet*; P*unc-122::dsRed*; *lin-15*] into *lin-15(n765)* mutants. The integrant was outcrossed with N2 three times and crossed into SJR1 to make SJR16 (used for recordings in Fig. 5 A-D).

Finally, for recording first- and second-layer interneurons simultaneously, *sjxIs8* was generated by integrating *sjxEx8*[P*npr-9::NLS-wrmScarlet*; P*ttx-3::mNeptune::GCaMP6s*; P*ceh-16::mNeptune::GCaMP6s*; P*gcy-28d::mCherry::GCaMP6s*; *lin-15*] into *lin-15(n765)* mutants. The integrant was outcrossed three times with N2 and crossed into ADS1001 to generate SJR15 (used for recordings in Fig. 4).

### Cultivation and microscopy

The animals were grown in a 20°C incubator on NGM plates seeded with OP50 bacteria. At the stage of young adult, they were transferred into a microfluidic polydimethylsiloxane (PDMS) arena for recording. The microfluidic chip was customized based on the design presented in ref. [47].

The recording was performed using a spinning disc confocal microscope (Nikon Eclipse Ti2 and Yokogawa CSUX1FW) and two Andor Zyla 4.2MP Plus cameras, one recording the green and the other recording the red channel. High-resolution images were collected through a 40× Nikon Plan Fluor Oil DIC N.A. 1.30 objective. Green (GCaMP6s) and red (mCherry, mNeptune, wrmScarlet) channel 3D volumetric stacks were obtained with an exposure time of 10 ms at approximately 3 volumes per second.

### Extraction of calcium activity

To extract the calcium activity of each neuron, we identified the 30% brightest red (mCherry, mNeptune, wrmScarlet) channel pixels in each neuronal volume. We then computed for these pixels the ratio (*R*(*t*)) of mean green (GCaMP6s) intensity to mean red intensity. Furthermore, to exclude the effects of outliers resulting from the poor annotation of frames, we dropped the second lowest and second highest percentile neuronal activities and smoothed the remaining recording tracks using a 1-D Gaussian filter with a standard deviation of 3.

Finally, the activity of each neuron at time *t* was computed using:

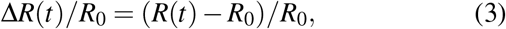

*R*_0_ is the lowest 1st percentile of *R*(*t*) in the recording.

## CODE AND DATA AVAILABILITY

The code and sample 4D datasets of pan-neuronal nuclear marked worms and multi-neuron cytosolic marked worms are available at https://github.com/lpbsscientist/targettrack.

## ACKNOWLEDGEMENTS

AD, KK, MBK, SJR were supported by the École polytechnique fédérale de Lausanne (EPFL), the Helmut-Horten Foundation, the Swiss Data Science Center grant C20-12, and an EPFL Interdisciplinary Seed Fund. We thank Nicholas Greensmith for help developing the GUI; Guillaume Obozinski for feedback; Albert Lin for help constructing strains; Matthieu Schmidt and Alice Gross for collecting and analyzing data.

## AUTHOR CONTRIBUTIONS

ADTS, CFP, KK, SJR conceived the project; CFP, KK, MBK, VS collected the data; CFP prepared the data, developed the neural network, suggested targeted augmentation, and implemented the method; MBK and CLJ adapted the method for 3D volumes; CFP, MBK ran the evaluations; CFP, MBK, AD developed the GUI; CFP, ADTS, SJR, MBK, CLJ wrote the manuscript; ADTS, SJR initiated and supervised the project.

## Supplemental figures

**Figure S1.**
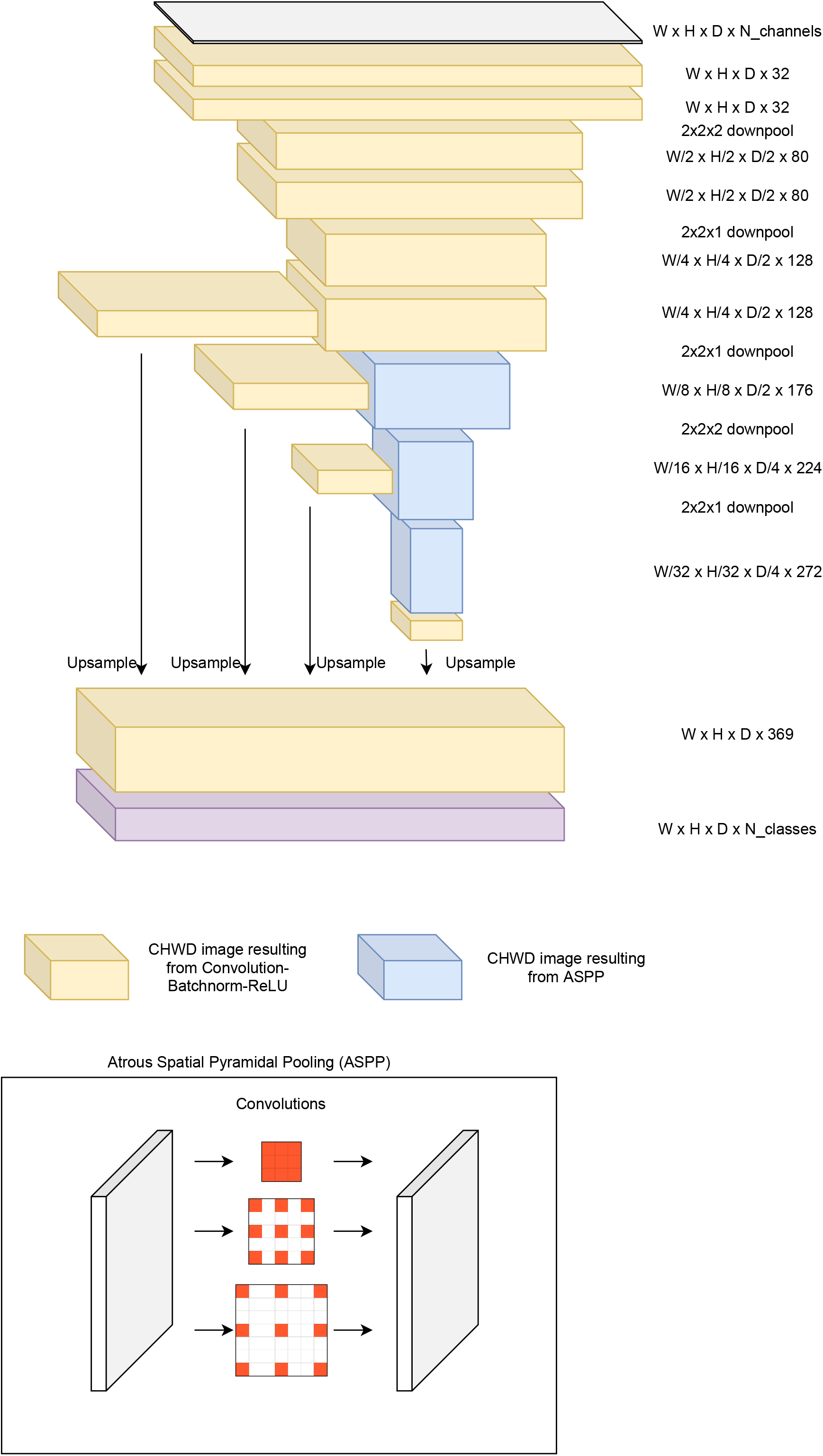
Illustration of the neural network architecture and of the atrous convolutions.

**Figure S2.**
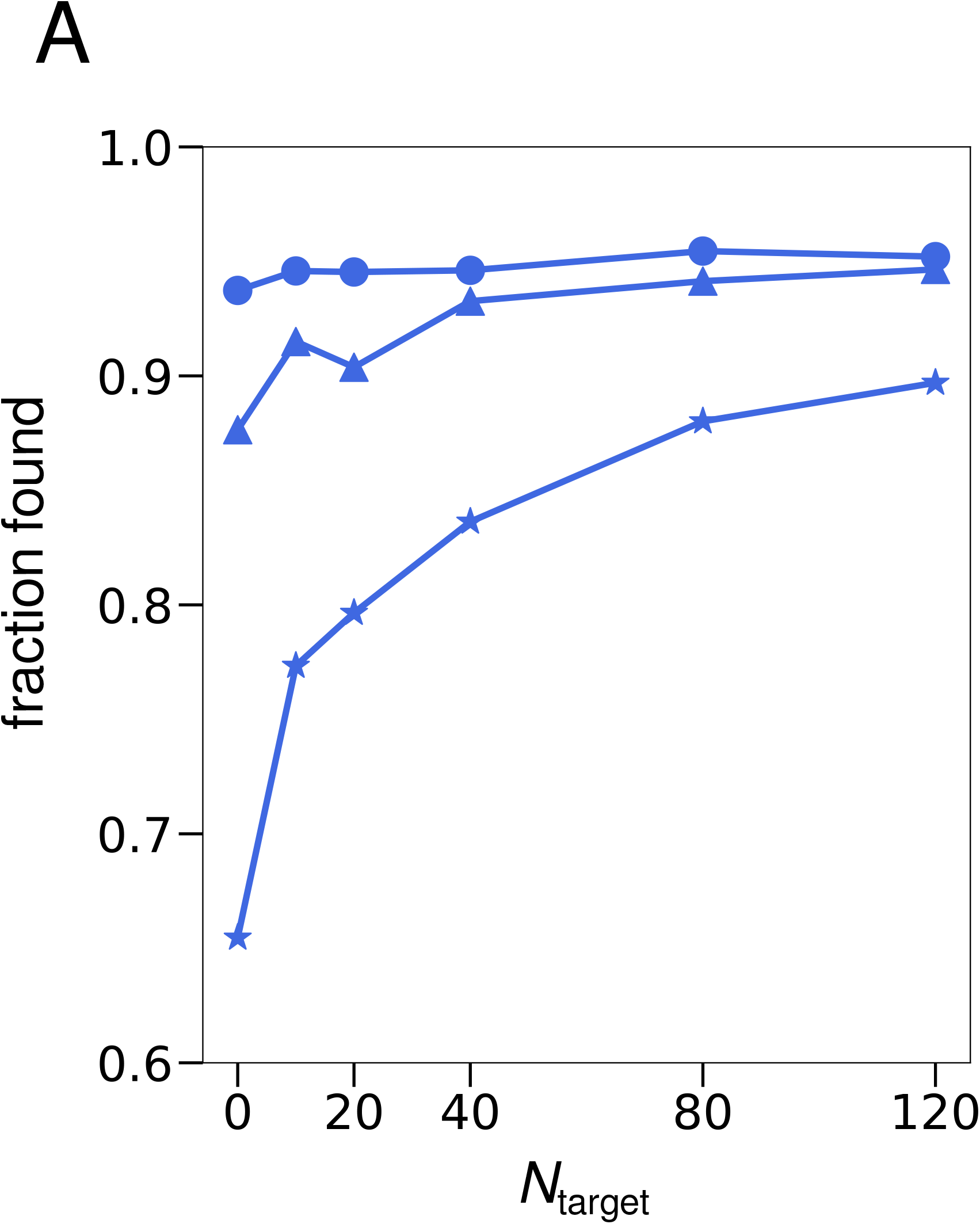
Fraction of objects found for one recording (blue plot in Fig. 4 A) with *N*_GT_ = 5 (stars), *N*_GT_ = 15 (triangles), or *N*_GT_ = 25 (circles) ground-truth annotations as a function of the target set size *N*_target_.

**Figure S3.**
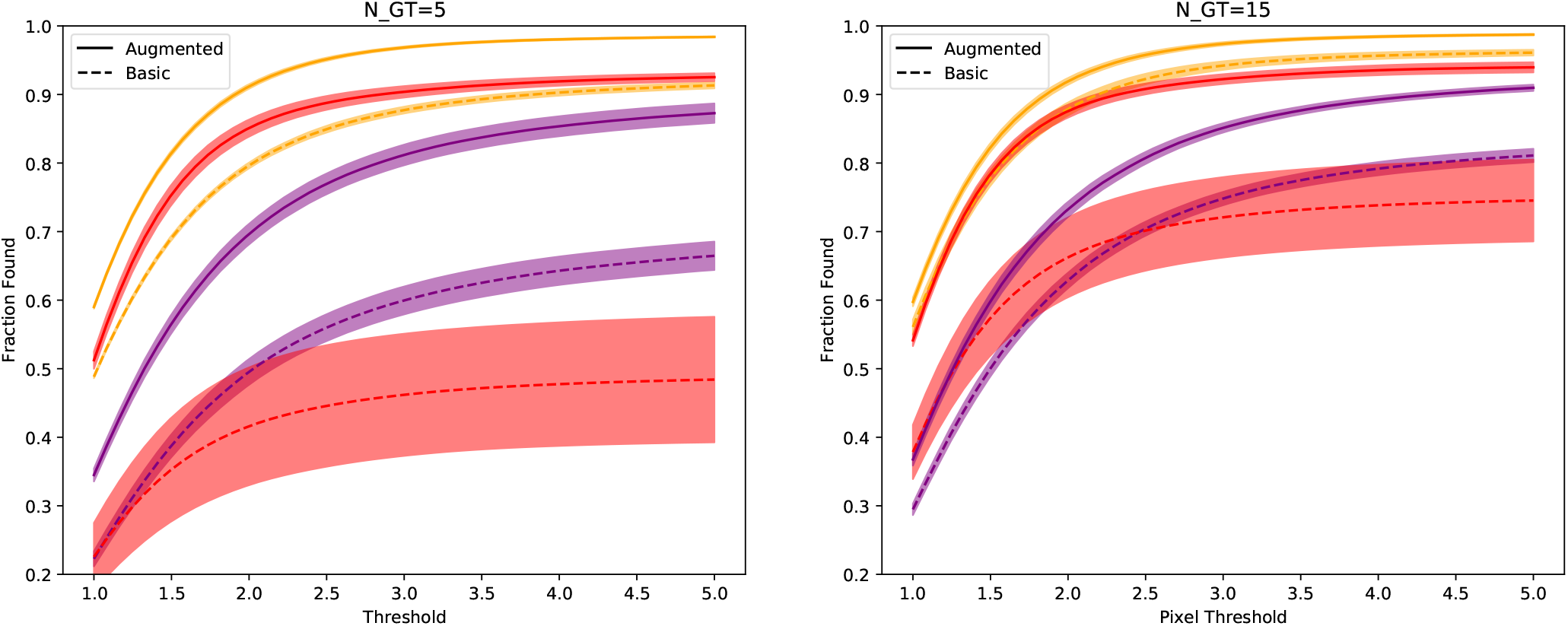
Fraction of key-points found depending on the pixel threshold for *N*_*GT*_ = 5 (left) and *N*_*GT*_ = 15 (right).

**Figure S4.**
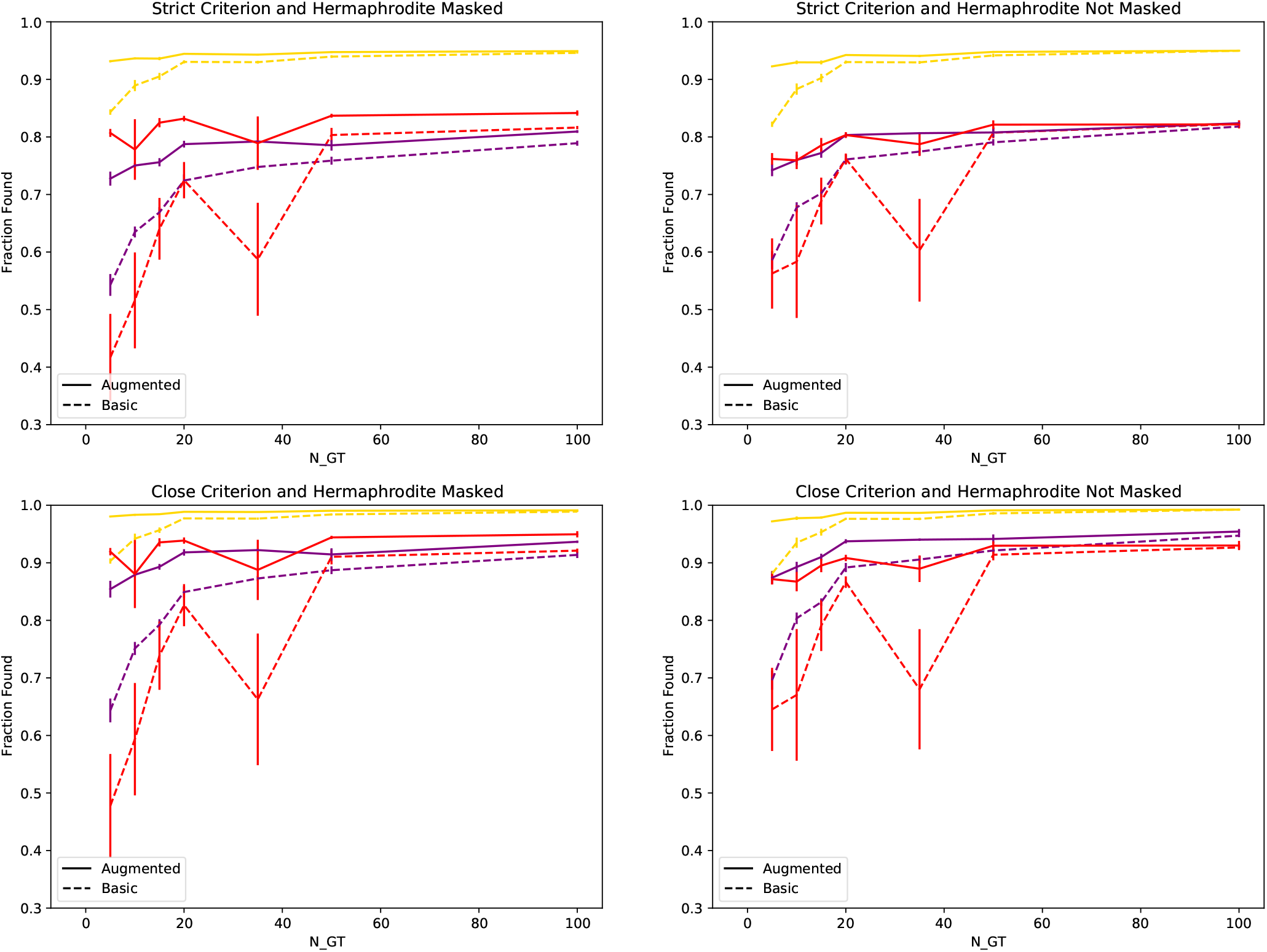
(top left) Fraction of key-points successfully found when the strict criterion is applied and the hermaphrodite is masked out (top right). Fraction of key-points successfully found when the strict criterion is applied and the hermaphrodite is not masked out (bottom left). Fraction of key-points successfully found when the strict criterion is not applied and the hermaphrodite is masked out (bottom right). Fraction of key-points successfully found when the strict criterion is not applied and the hermaphrodite is not masked out.

**Figure S5.**
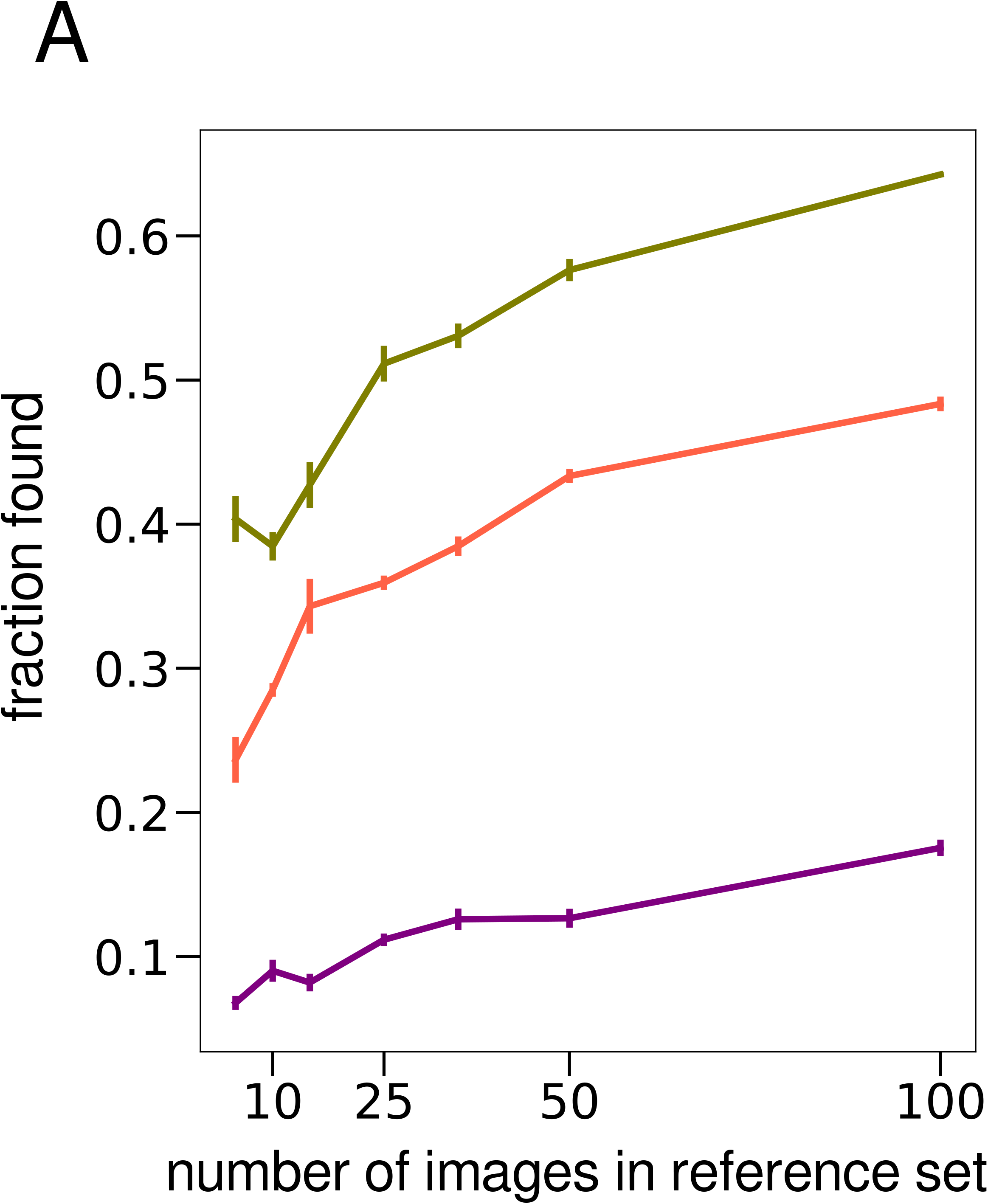
Fraction found for the CPD method.

## Supplemental note 1: Details of the CPD method

We use CPD (Coherent Point Drift) together with a nearest neighbor matching scheme as a simple tracking method for comparison. We assume a perfectly solved segmentation problem using the ground truth pointset from manual annotation. The algorithm is as following:

1. We begin with N annotated pointsets, i.e., points are provided with labels.
2. For each non-annotated pointset:
  a. The nearest annotated pointset among the N pointsets is found.
  b. The nearest annotated pointset found is deformed into the current pointset using CPD.
  c. The label of each point is assigned to be the label of the nearest deformed annotated pointset.

